# How did UGA codon translation as tryptophan evolve in certain ciliates? A critique of Kachale et al. 2023 *Nature*

**DOI:** 10.1101/2023.10.09.561518

**Authors:** Estienne Carl Swart, Christiane Emmerich, Kwee Boon Brandon Seah, Minakshi Singh, Yekaterina Shulgina, Aditi Singh

## Abstract

Ciliates are a widespread clade of microbial eukaryotes with the greatest diversity of nuclear genetic codes (at least eight) following a recent addition^1^. All non-standard ciliate genetic codes involve stop codon reassignments^1,2,3^. Two of these codes are ambiguous^1–3^, with “stop” codons either translated or terminating translation depending on their context^2,3^. Ambiguous genetic codes have arisen not only in ciliates, but also independently in trypanosomatids from the genus *Blastocrithidia*^4^ and an alveolate species from the genus *Amoebophrya*^5^. Two ambiguous genetic codes in ciliates share translation of UGA “stop” codons as tryptophan with *Blastocrithidia* and the *Amoebophrya* species. tRNA genes with complementary anticodons to reassigned UAA and UAG stop codons have invariably been found in ciliate species that translate these codons^1,2^. Furthermore, though a UGA-cognate tRNA^Cys^_UCA_ was reported in *Euplotes*^6^, a ciliate genus that translates UGA as cysteine, vexingly, no nuclear genome-encoded tRNA^Trp^_UCA_ has been found in ciliate species with UGA tryptophan codons. Recently, Kachale et al. provided evidence for UGA translation as tryptophan in *Blastocrithidia nonstop* and the ciliate *Condylostoma magnum* using 4 base pair anticodon stem (AS) near-cognate tryptophan tRNA^Trp^_CCA_’s, rather than the typical 5 base pair stem tRNAs^7^. New tRNA data we report from additional ciliates bolsters this hypothesis. Kachale et al. also hypothesised that a particular amino acid substitution in the key stop codon recognition protein, eRF1 (eukaryotic Release Factor 1), favours translation of UGA as tryptophan instead of termination^7^. Contrary to Kachale et al, we propose such substitutions favouring reduced eRF1 competition enhancing “stop” codon translation do not need to occur concomitantly with tRNA alterations or acquisitions to evolve new genetic codes via stop codon reassignment. We report multiple instances of the substitution investigated in Kachale et al. 2023 that have not led to UGA translation, and multiple ciliate species with UGA tryptophan translation but without the substitution, indicating it is not necessary. Consistent with the ambiguous intermediate hypothesis for genetic code evolution, experimental evidence and our observations suggest continued potential ciliate eRF1-tRNA competition.

## Critique

There are two important issues in Kachale et al. 2023 to resolve from the outset, both clarified in Supporting Information. One: a reported 5 bp AS tryptophan tRNA (in their Fig. 3a) which did not lead to efficient stop codon readthrough^7^, likely originates from a bacterium present in the ciliate culture, not from *Condylostoma magnum*. Thus, *Condylostoma* probably only has nuclear genome-encoded 4 bp AS tRNA^Trp^_CCA_’s, further supporting the proposal that they are necessary for efficient UGA translation as tryptophan. Two: the authors express uncertainty about the genetic codes used by several ciliate species, particularly relating to UGA codons and tRNAs with potential complementary anticodons; this can partly be traced to incorrect annotations in reference databases or earlier publications, that have since been superseded. This is pertinent to the interpretation of eRF1 substitutions and their role in UGA translation.

With their diverse genetic codes, ciliates are ideal for exploring hypotheses about these codes. Consistent with Kachale et al’s 4 bp AS tryptophan tRNA proposal for translation of UGA as tryptophan, we find such tRNAs occur not only in *Condylostoma magnum* (class Heterotrichea; UAR=Q/*, UGA=W/*)^2,3^ but also in another ciliate species with an ambiguous genetic code, *Loxodes magnus* (class Karyorelictea; UAG=Q, UAA=Q/*? [UAA may also be a stop], UGA=W/*)^8^, and in the ciliate genus *Blepharisma* (class Heterotrichea; Fig. 1), which appears to translate UGA unambiguously as tryptophan (UAR=*, UGA=W)^8^. A draft *Loxodes magnus* somatic genome assembly has over one hundred 4 bp AS tRNA^Trp^ genes but just six 5 bp AS tRNA^Trp^ genes (Fig. 1). The 5 bp AS tRNAs of *L. magnus* shown in Fig 1. have one or two unpaired T-stem bases and are co-located on a contig with similar sequences and secondary structures, particularly the T-stem, but different anticodons (CUU, CUA, CCU) corresponding to non-tryptophan codons. This suggests relaxation of selection, and that these are likely pseudogenes. In contrast, two other heterotrich ciliates with standard genetic codes^9,10^, *Stentor coeruleus* and *Fabrea salina*, only have 5 bp AS tRNA^Trp^ genes (Fig. 1). Among members of the *Paramecium aurelia* complex, which are not known to translate UGA as tryptophan (code: UAR=Q, UGA=*)^2^, we found a ciliate species, *Paramecium biaurelia*, with at least one 4 bp AS tRNA (Extended Data Fig. 1a) among its eight tRNA^Trp^ genes. For such tRNAs and additional ones in other organisms that are not evident pseudogenes, the ability to translate UGA as tryptophan should be carefully experimentally investigated in future.

**Fig. 1.**
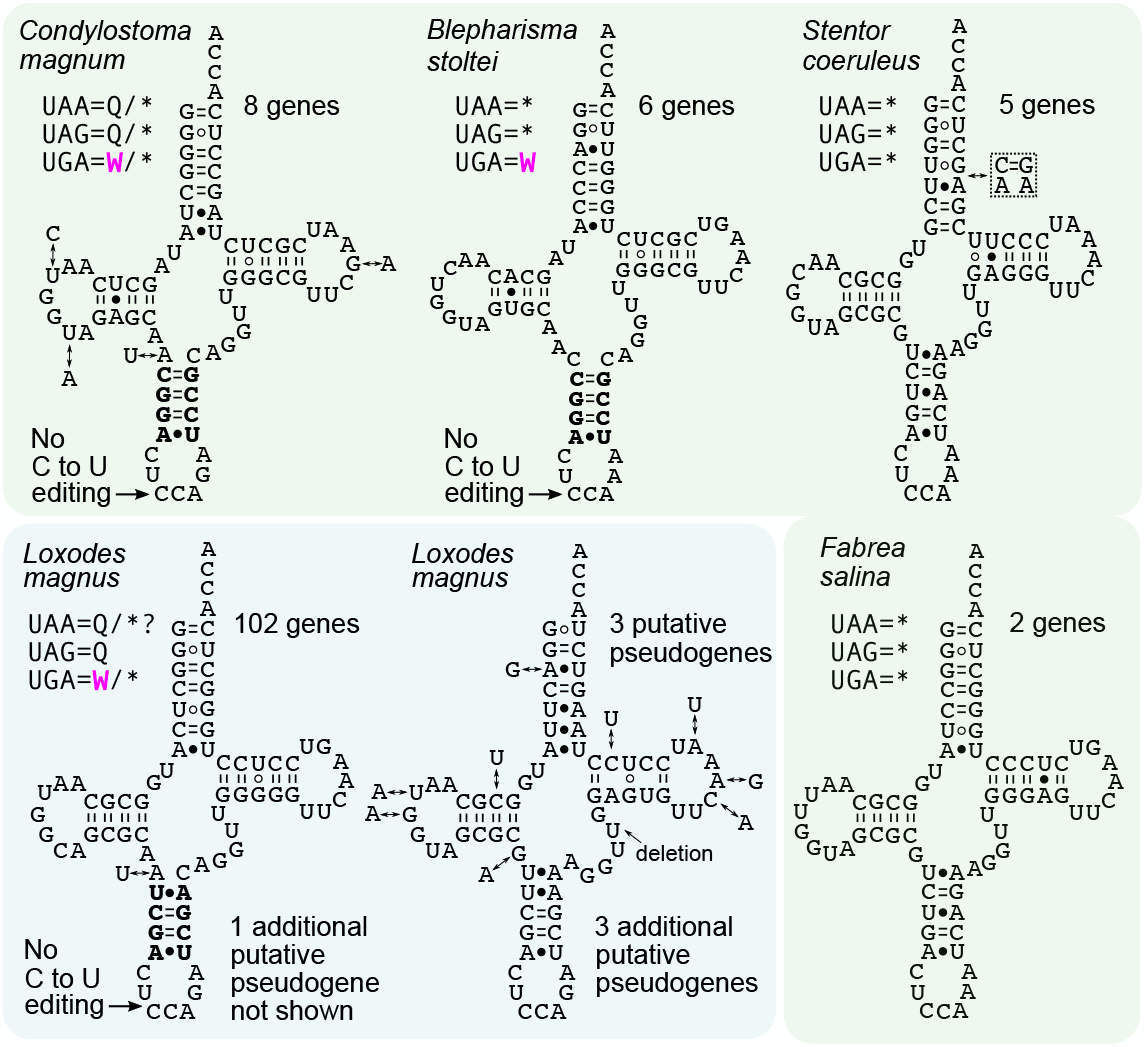
Predicted tryptophan tRNAs encoded in the macronuclear genomes of heterotrich and karyorelict ciliates. The two ciliate classes are indicated by different background colours: Heterotrichea - green; Karyorelictea - cyan. Nucleotide substitutions that differ between tRNA genes are indicated by double arrows. Stop codon reassignments are given under the species names. See Extended Data Fig. 1 for *Blepharisma*’s tRNA_UCA_ predictions (selenocysteine and mitochondrial). *Blepharisma japonicum* and *Blepharisma undulans* tRNA^Trp^_CCA_ also have 4 bp anticodon stems (Source Data Fig. 1 and Extended Data Fig. 1).

In a human pathogenic trypanosomatid *Leishmania* species, tRNA C-to-U editing of the wobble (5’) anticodon base of nuclear genome-encoded mitochondrial tRNA^Trp^_CCA_’s generates UCA anticodons that enable mitochondrial UGA codon translation^11^. Kachale et al. reported no editing of cytosolic *Blastocrithidia* tRNA^Trp^_CCA_’s^7^ that could permit translation of cytosolic UGAs. Previously, tRNA sequencing did not reveal C-to-U editing of the *Condylostoma magnum* tRNA^Trp^_CCA_ anticodon wobble base that would generate an anticodon complementary to UGA codons^2^. We have also observed no appreciable C-to-U editing (> 0.1% of tryptophan tRNAs) in *Blepharisma* and *Loxodes* 4 bp AS tRNA^Trp^ sequences (in 172,929 and 9,721 unique reads, respectively; Supplementary Table 1, Source Data Fig. 1).

Anticipating the discovery of natural 4 bp AS tRNA^Trp^_CCA_’s in *Blastocrithidia* and the ciliates *Condylostoma, Blepharisma*, and *Loxodes*, the idea that tRNA AS mutations can enhance decoding of near-cognate codons was previously explored in back-to-back papers by Schultz and Yarus^12,13^. In the first paper, by extensive mutational screening of *Escherichia coli* tRNA su7 G36, a derivative of a tryptophan suppressor tRNA with a CUG anticodon, they found that mutations that disrupt the top AS stem base pair — creating a 4 bp AS stem — led to the most translation of UAG^12^, which involves G-U wobble pairing at the 1^st^ codon position that is normally disallowed. In the second paper, using su7 tRNA_CUA_ they showed that mutations the disrupt the top AS stem base pair promote UAA translation^13^, which involves mismatched 3^rd^ position A-C like that required for UGA translation by 4bp AS tRNA_CCA_’s^7^.

Particular amino acid substitutions in homologs of the protein that recognizes stop codons, eRF1, were formerly thought to be associated with loss of stop codon recognition necessary for the evolution of particular genetic codes in ciliates^14,15^. With the benefit of additional eRF1 sequences and ciliate genetic codes, we previously reported multiple counterexamples to such associations^2^. Kachale et al. proposed that a single amino acid substitution in eRF1, from Ser67 to Ala/Gly67 (numbered with respect to yeast eRF1), may be needed for loss of UGA termination in conjunction with a shorter tRNA^Trp^ anticodon stem for efficient UGA translation as tryptophan^7^. However, this substitution is present in eRF1’s of multiple ciliates that use the standard genetic code: *Stentor coeruleus, Fabrea salina* and *Climacostomum virens* (all members of class Heterotrichea; like *Condylostoma* and *Blepharisma*; Fig. 2a). eRF1 of the ciliate *Pseudocohnilembus persalinus* (class Oligohymenophorea), which has the genetic code UAR=Q and UGA=*, also has Ala67, and so too does eRF1 of the diplomonad flagellate *Giardia intestinalis* (standard genetic code).

**Fig. 2.**
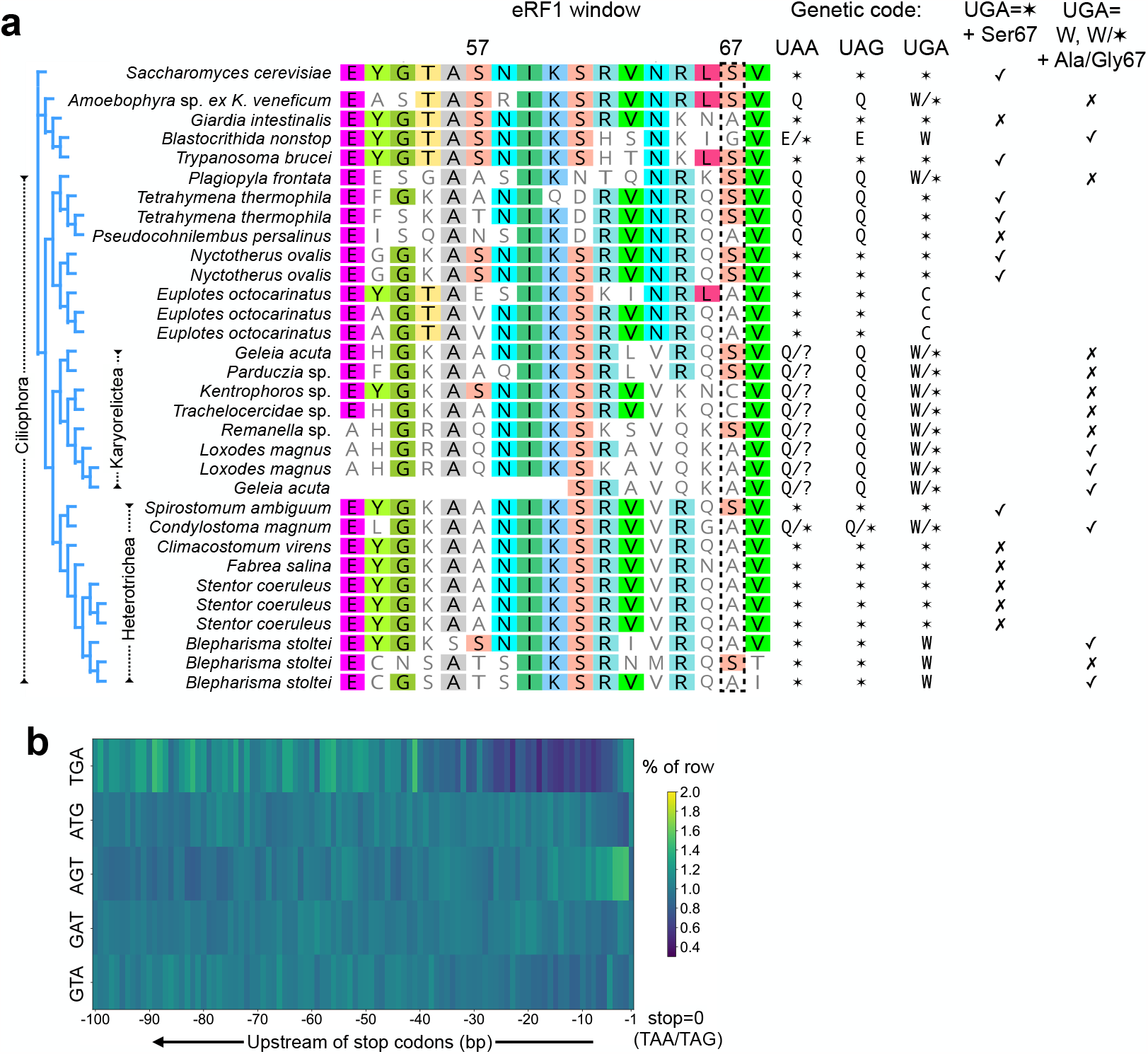
eRF1 substitutions and potential signals of eRF1-tRNA competition in *Blepharisma*. (a) eRF1 coordinates are given according to that of *Saccharomyces cerevisiae*, and the alignment window is the same as that in Kachale et al. 2023. The complete eRF1 alignment along with the sources of the sequences is provided in Source Data Fig. 2. For the genetic codes of each species, stars indicate stop codons, and question marks indicate possible stop codons. Check marks and crosses respectively indicate agreements and disagreements with respect to the proposed UGA assignment/eRF1 substitution at position 67. The eRF1 phylogeny to the left was generated by RAxML. The second *Geleia acuta* eRF1 paralog is encoded by an incomplete transcript. (b) Codon frequency upstream of predicted *B. stoltei* stop codons for the permutations of “T”, “G” and “A” bases. For the complete codon frequency matrix, see Extended Data Fig. 2a.

Furthermore, all karyorelict ciliates translate UGA as tryptophan within mRNA coding sequences (and use it as a stop at the ends of coding sequences)^8^. One species, *Loxodes magnus*, has eRF1’s with Ala67, but other species’ eRF1’s have either Ser67 or Cys67 (Fig. 2a). Recently the ciliate species *Plagiopyla frontata* (class Plagiopylea) was reported to have ambiguous UGA codons that are translated as tryptophan in coding sequences (genetic code UAR=Q, UGA=W/*)^1^. eRF1 from this distantly related ciliate has Ser67. So too does eRF1 from the alveolate *Amoebophrya* sp. ex *Karlodinium venificum*. Thus, the Ala/Gly67 substitution is present in ciliate species without UGA translation and is not necessary in multiple ciliate species which translate UGA as tryptophan. Ala/Gly67 also appears unnecessary in the *Amoebophrya* species.

Unlike bacteria which have two proteins that recognize two stop codons each, RF1 and RF2, standard genetic code model eukaryotes, like yeast, typically have a single “omnipotent” protein, eRF1, that recognizes all three stop codons. eRF1 paralogs were previously noted to have arisen independently in certain ciliate genera, including *Euplotes* and *Tetrahymena*^16,17,18^. The frequent occurrence of such paralogs in ciliates raises the possibility some may have subfunctionalized, like RF1 and RF2, with different stop codon recognition capabilities, but this needs experimental determination. *Blepharisma stoltei* has three divergent eRF1 paralogs (60-72% amino acid identity for the three pairwise comparisons), of which the most highly transcribed one (BSTOLATCC_MAC3627; mean 540 RPKM, standard deviation 50 RPKM for a developmental time series^19^) has Ala67, but there is also an eRF1 paralog with low transcription (mean 9.9 RPKM, standard deviation 6.2 RPKM) and Ser67 (Fig. 2a). Gene expression of the more divergent *Blepharisma* eRF1 paralogs is comparable to that of the ancient eRF1 paralog Dom34/Pelota (BSTOLATCC_MAC12938; mean 6.8 RPKM; standard deviation 2.3 RPKM), a protein responsible for the translation-associated process “No-Go decay”^16^. It is conceivable that these paralogs have functionally diverged, now serving an alternative role like Dom34/Pelota.

Though we have only observed UGA codons translated as tryptophan in *Blepharisma stoltei, in vitro* translation experiments suggest *Blepharisma japonicum*’s ortholog of the highly transcribed *B. stoltei* eRF1, also with Ala67, can recognize all three stop codons, but UGA the most weakly^20^. Correspondingly, with some capacity of *Blepharisma* eRF1 to recognize UGA as stop codons, a signal of potential competition between *B. stoltei* eRF1 and tRNA^Trp^ can be observed in the form of UGA codon depletion in a region beginning 25-30 codons upstream of UAR stop codons (Fig. 2b). A similar depletion was observed in the karyorelict ciliates and the heterotrich ciliate *Condylostoma* which do use UGA as a stop close to transcript ends^2,8^. Interestingly, depletion of UAA and UGA codons occurs in a similar region before stops in *Blastocrithidia nonstop*, contrasting with the constancy of reassigned UAG codons^7^. This suggests eRF1-tRNA competition not only for UAA but also for UGA in this species.

While the type of amino acid substitution proposed by Kachale et al. may certainly substantially enhance translation, it should be noted that such substitutions are not a prerequisite for the acquisition of a new genetic code under the hypothesis which best fits the evolution of the ambiguous stop/sense genetic codes, the “ambiguous intermediate hypothesis”^21^. Instead, in a transitional evolutionary phase, codons may be interpreted in two ways, with potential eRF1-tRNA competition. With time, beneficial mutations or modifications in either the tRNA or eRF1 (or other components of translation) that reduce competition may be selected.

Instead of focusing on individual eRF1 substitutions, we propose future investigations should more generally explore the structure of non-standard genetic code eRF1’s captured in translation termination in the context of their own ribosomes. New genetic codes involving stop codon reassignments have had ample opportunity to evolve in eukaryotes through a combination of tRNA and eRF1 mutations, but are limited to just a few clades, most notably having radiated in ciliates. We thus infer that an additional aspect has enabled genetic code evolution in these prolific microbes, and continue to suggest that this may be their ability to either tolerate or resolve genetic code ambiguity.

## Contributions

Conceptualization: E.C.S. Investigation: E.C.S., B.K.B.S, M.S. Methodology: E.C.S, C.E., B.K.B.S, A.S. Resources: all authors. Writing: all authors. Supervision: E.C.S.

## Figure captions

**Extended Data Fig. 1.**
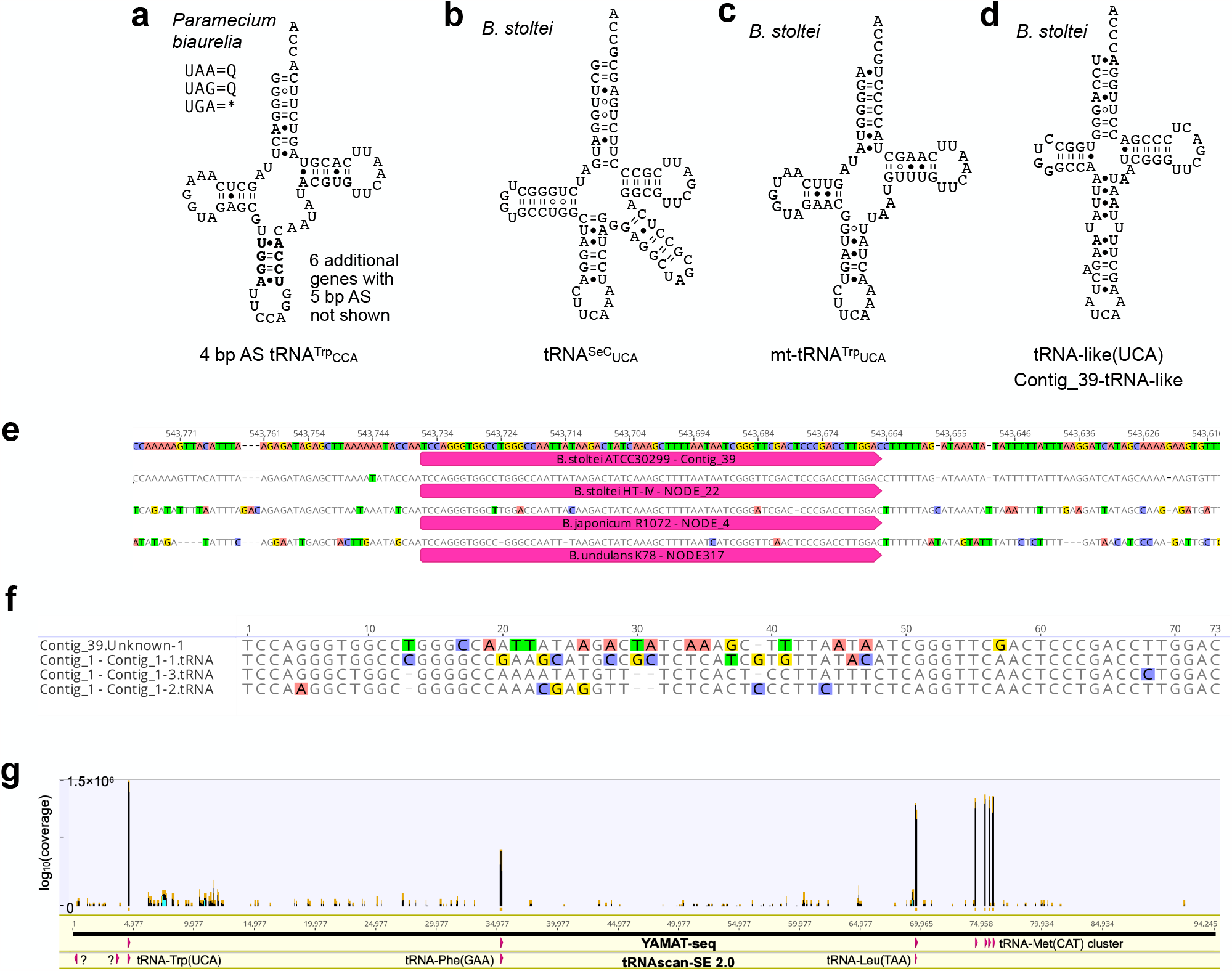
Secondary structures of *Paramecium biaurelia* 4 bp AS tRNA^Trp^_CCA_ and *Blepharisma* tRNA_UCA_’s and a tRNA-like molecule with a possible UCA anticodon. (a) *Paramecium biaurelia* tRNA^Trp^_CCA_; terminal nucleotides predicted by tRNAscan-SE 2.0. (b) *B. stoltei* tRNA^Sec^_UCA_. (c) *B. stoltei* mitochondrial tRNA^Trp^_UCA_. (d) tRNA-like molecule with potential UCA anticodon in *B. stoltei*. (e) Multiple sequence alignment of tRNA-like sequences from *Blepharisma* spp. (f) Multiple sequence alignment of paralogs of tRNA-like sequence paralogs in *B. stoltei* ATCC30299. (g) YAMAT-seq mapping to the *B. stoltei* ATCC30299 mitochondrial genome.

**Extended Data Fig. 2.**
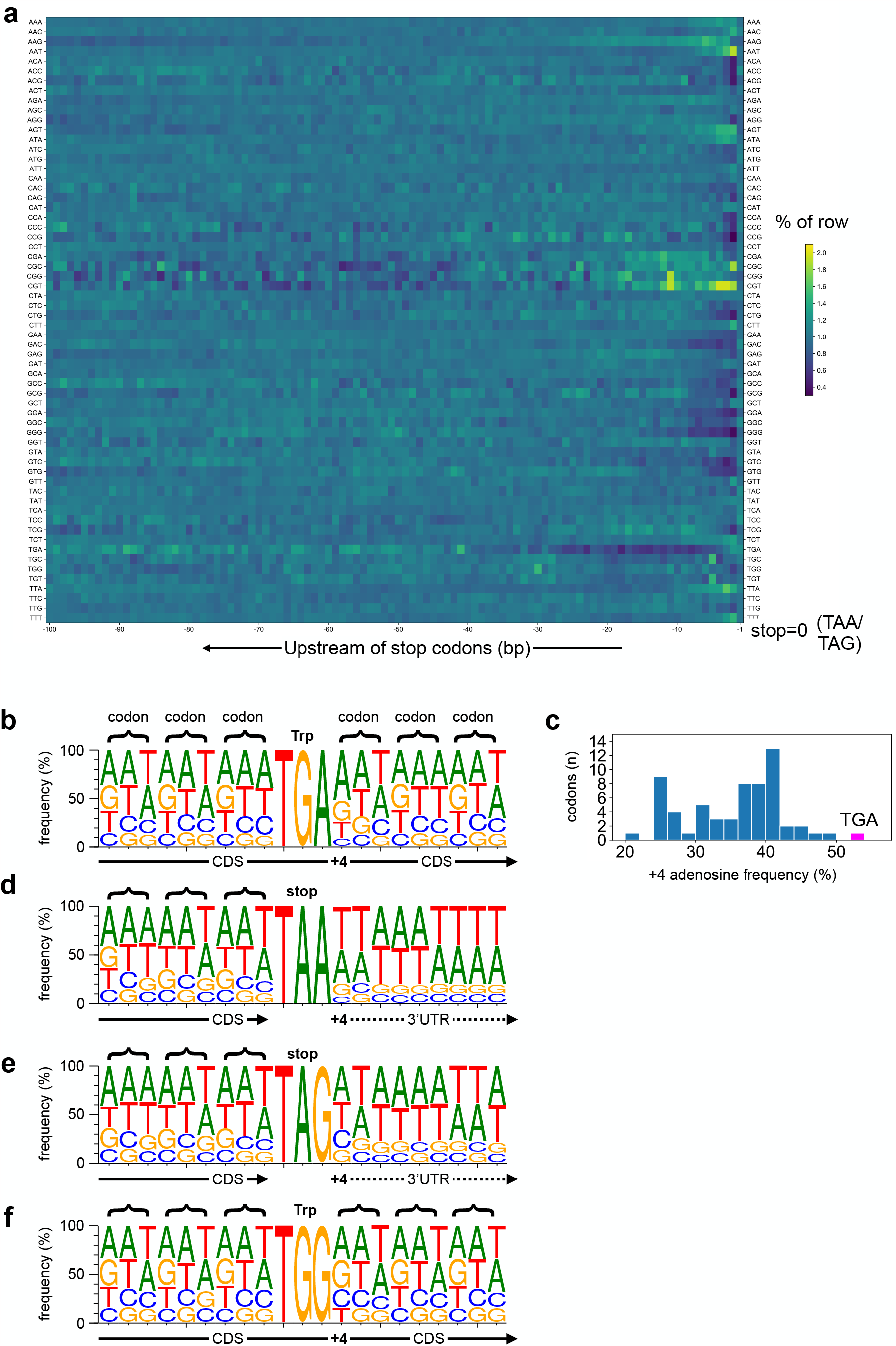
Codon usage before stops and base frequencies around stops. (a) Codon frequency upstream of predicted *B. stoltei* stop codons. (b) Base frequencies flanking *B. stoltei* TGA codons (n=44087). (c) Adenosine frequency of +4 at base immediately downstream of translated *B. stoltei* codons. (d) Base frequencies around TAA (n=21002) stop codons. (e) Base frequencies around TAG (n=4707) stop codons. (f) Base frequencies around TGG tryptophan codons (n=78535).

