## Supplemental Information for "How did UGA codon translation as tryptophan evolve in certain ciliates? A critique of Kachale et al. 2023 *Nature*"

#### Evidence that a 5 bp anticodon stem tryptophan tRNA reported for *Condylostoma* in Kachale et al. 2023 is a bacterial contaminant

In Fig 3a from Kachale et al. 2023, the 5 bp AS $tRNA^{Trp}_{CCA}$ reported for *Condylostoma magnum* is encoded on a 11.7 kb contig from DNA we previously isolated, sequenced and assembled (ENA: CVLX01007640.1). At 41% GC, the base composition of this contig is substantially higher than the typical *Condylostoma* somatic genome base composition (*Condylostoma* transcripts have a mode of 33% GC; the genome has more AT-rich intergenic regions[1]). Such contigs were not used for analyses of the *Condylostoma* genetic code and subsequent analyses, which is why only 4 bp AS $tRNA^{Trp}_{CCA}$'s were previously reported as being present in the macronuclear genome[1]. Top matches of BLASTX searches using sequences downstream of the tRNA with the standard genetic code versus GenBank's NR database (update date: 12 January 2023) were bacterial. These matches also lack the frequent in-frame stops expected if they used *Condylostoma*'s alternative code. Since the *Condylostoma* culture from which we isolated the DNA was not axenic, it appears this tRNA originates from a contaminating bacterium. Therefore, *Condylostoma* probably has only 4 bp AS $tRNA^{Trp}_{CCA}$'s.

#### Clarification of the correct genetic codes and tRNAs cognate to UGA reported in Kachale et al. 2023

Kachale et al. 2023 contains three different figures with different subsets of eRF1 sequences: their Fig. 4a; Extended Data Fig. 10; and an external data figure from Figshare (https://figshare.com/projects/tRNA_anticodon_stem_length_variations_are_critical_for_stop_codon_reassignment/129167 - Analysis of eRF1, UPF1/2 and tRNAs in ciliate genomes). The first figure only shows *Condylostoma magnum* eRF1, with alanine at the position of interest (67); the second figure shows additional ciliate eRF1s; the last figure focuses on and shows the widest sampling of ciliate eRF1s and reports stop codon reassignments. We focus on the last figure, as it has the strongest bearing on the hypothesis for the necessity of the eRF1 Ser67 to Ala/Gly67 in permitting UGA translation as tryptophan.

In Kachale et al.'s external data figure, the correct stop codon assignments are reported for the heterotrichs *Fabrea salina* and *Climacostomum virens*, based on codes we previously predicted[1,2], but, before them as alternatives are listed incorrect reassignments to cysteine from the National Center for Biotechnology Information (NCBI) "The Genetic Codes"[3], part of the NCBI Taxonomy Database (accessed 17 March 2023)[4]. Correct assignments of UGA codons as stop codons were also independently determined and reported for *F. salina*[5,6]. *Stentor coeruleus* uses the standard genetic code[7], which means it does not translate UGA as cysteine as reported by Kachale et al. 2023.

We checked predicted *Stentor coeruleus* tRNAs from ARAGORN and tRNAscan-SE 2.0; besides the usual selenocysteine tRNAs ($tRNA^{Sec}_{UCA}$'s), we found two tRNAs with UCA anticodons complementary to UGA, one of which may be a pseudogene with two successive non-canonical A-G base-pairs present in the predicted anticodon stem (GenBank accession MPUH01000092.1: bases 29518 to 29590; the other tRNA is encoded by the *Stentor* mitochondrial genome[7] (GenBank accession MPUH01000652.1: bases 9888 to 9960). So, the symbol indicated in the external data figure for a nuclear genome-encoded $tRNA_{UCA}$, should be a "-", not a "+". Similarly, we deduce that the $tRNA^{Trp}_{UCA}$ indicated as "+" for *Condylostoma magnum* is either a mitochondrial genome-encoded $tRNA^{Trp}$ (since the mitochondrial genome was co-assembled with the nuclear genome) or a $tRNA^{Sec}_{UCA}$.

Two genetic codes are listed by NCBI's taxonomy for different *Nyctotherus* species (codes 6 and 10), but the most thorough investigation of the *Nycotherus ovalis* MAC genome reports that it uses the standard genetic code (code 1)[8]. There is no evidence to suggest this species translates UGA as cysteine (code 10) as reported in Kachale et al. 2023's external data figure. Therefore, all the NCBI genetic codes listed in the external data figure are incorrect. A summary of incorrect NCBI genetic codes and what they currently ought to be is given in Supplementary Table 2.

### Clarification of widespread confusion in the specification of the *Blepharisma* genetic code

Kachale et al. 2023 report the correct genetic code, to the best of our knowledge, for *Blepharisma japonicum*. However, incorrect *Blepharisma* genetic codes persist in multiple websites on the Internet and in bioinformatics software, and so we wish to alert potential genetic code investigators and users of such software of the issue.

Based on analyses of stop codon recognition capabilities of the release factor eRF1 the species *Blepharisma japonicum* was once thought likely to use the standard genetic code[9]. Due to the limited sequence data available and fairly low frequency of UGA codons, the reassignment of UGA as tryptophan was not detected. This reassignment was reported a decade prior for *Blepharisma americanum*[10], and subsequently verified for a *Blepharisma japonicum* transcriptome assembly[1,6]. We found the same genetic code (UGA=W; UAR=*) in *Blepharisma stoltei*[11], and so we think it likely that this code is used throughout the genus *Blepharisma*.

"The genetic codes"[3] from NCBI underpins translation of protein-coding sequences of all the major sequence databases (i.e. GenBank, The European Nucleotide Archive, DNA Data Bank of Japan), and this resource is frequently referenced during investigations of genetic code evolution. So, it is important it be kept up-to-date and corrected. Confusion in the literature about what genetic code *Blepharisma* uses has persisted for many years due to an error in these codes (dating back to at least 2003)[12]. The incorrect genetic code (number 15) differs from the standard genetic code in the reassignment of the stop UAG to glutamine, but the paper cited in support of this made no such claim[13]. In more recent versions of NCBI's genetic codes, number 15 has been removed, but prior to this the error spread through to bioinformatics software that uses the NCBI genetic codes. Among others this currently (8 March 2023) includes: software routinely used in gene prediction — AUGUSTUS[14], the general-purpose genomic sequence manipulation suite — Geneious[15], and bioinformatics programming libraries like BioPython[16].

To translate *Blepharisma* genes, it is currently possible to use NCBI genetic code 4 ("The Mold, Protozoan, and Coelenterate Mitochondrial Code and the Mycoplasma/Spiroplasma Code"). However, due caution should be exercised because this code has numerous alternative initiation codons other than AUG, for which there is currently no evidence in *Blepharisma*. We have not yet succeeded in convincing the maintainers of NCBI's "The Genetic Codes" to reinstate the old code number (15) together with the correct code.

#### Clarification of eRF1 present in *Amoebophrya* sp. *ex Karlodinium venificum*

Two eRF1 proteins are shown for *Amoebophrya* sp. *ex Karlodinium venificum* in Kachale et al. 2023's Extended Data Fig. 10. This is a parasitic species hosted by the dinoflagellate *Karlodinium veneficum*. Using translated BLAST searches with *Homo sapiens* eRF1 as the query sequence, we searched the two transcriptomes, GenBank accession GGWB00000000 for the parasite, and GWG00000000 for the host, separated out from a common RNA-seq assembly of this parasite and its host[17]. We found only one eRF1 protein (Fig. 2a) with Ser67 in the parasite transcriptome (encoded by two transcript isoforms, GGWB01013280.1 and GGWB01013281.1). Interestingly, none of this protein's 14 glutamine codons is UAA or UAG, and none of its 6 tryptophan codons is UGA, i.e. its gene can be correctly translated using the standard genetic code. The other eRF1 (Gly67) reported by Kachale et al. 2023 is encoded by a transcript that was classified as belonging to the dinoflagellate host, *Karlodinium venificum*, by the investigator who deposited the transcriptomes, since it is more GC rich (47%) than typical parasite transcripts (35% GC).

#### Investigation of *Blepharisma stoltei* tRNAs and tRNA-like molecules

The draft *Blepharisma stoltei* MAC genome[11] is currently (13 March 2023) the most complete one of any ciliate that translates UGA as tryptophan. In this genome, 169 and 176 tRNAs were predicted by tRNAscan-SE 2.0[18] and ARAGORN[19], respectively, with the predictions largely overlapping (Source Data Fig. 1). The 49 distinct anticodons of the tRNAs predicted by tRNAscan-SE are capable of translating all 64 codons, as well as methionine start codons and selenocysteine codons.
In the *Blepharisma stoltei* MAC genome, neither tRNAscan-SE 2.0 nor ARAGORN predicted any tRNAs with anticodons cognate to UGA other than a typical selenocysteine tRNA (87 bp long;

Extended Data Fig. 1b), which has a characteristic long variable arm. Like other ciliates, *Blepharisma*'s MAC genome has clear selenoprotein genes, such as a glutathione peroxidase (BSTOLATCC_MAC25242). In such cases, gene prediction software incorrectly translated UGA codons as tryptophan, instead of selenocysteine. The 3' untranslated regions (UTRs) of the genes encoding these proteins are also longer than usual and have potential SECIS element stem-loop structures, necessary for selenocysteine incorporation (e.g. predicted by RNAfold[20]). As in other ciliate mitochondrial genomes which translate UGA as tryptophan[21], tRNAscan-SE 2.0 also predicted a $tRNA^{Trp}_{UCA}$ in the *B. stoltei* ATCC30299 mitochondrial genome (Extended Data Fig. 1c).

YAMAT-seq is capable of identifying potential tRNAs that tRNAscan-SE 2.0 does not predict with current parameters, such as those in the *B. stoltei* ATCC30299 mitochondrial genome (Extended Data Fig. 2f). In particular, though tRNAscan-SE 2.0 did not detect mitochondrial $tRNA^{Met}_{CAU}$ genes in the *Blepharisma* mitochondrial genome, despite trying a couple other mitochondrion specific parameters (switches: "-M vert" and "-M mammal") in addition to the default, eukaryotic prediction parameters, the mapped YAMAT-seq reads allowed us to identify a cluster of four methionine tRNA genes encoded by the mitochondrial genome. With the "-mt" mitochondrial switch these tRNA genes were predicted by ARAGORN, but so too were additional tRNA genes that lack YAMAT-seq support.

We identified six loci in the *Blepharisma* MAC genome lacking a tRNA predicted by tRNAscanSE 2.0 but with high coverage of YAMAT-seq reads mapped by HISAT2[22], i.e. 620x-7800x, vs. a median of 5500x for tRNAs predicted by tRNAscan-SE 2.0. RNA secondary structures predicted from one of these loci by RNAfold[20] contained both a cloverleaf structure and a potential UCA anticodon (Contig_39-tRNA-like(UCA); 7800x coverage; Extended Data Fig. 1d). This tRNA-like molecule does not resemble *Blepharisma*'s $tRNA^{Trp}_{CCA}$'s (Fig. 1). Three additional putative tRNA-like molecules (7700x coverage) clustered together on Contig_1 between a pair of protein-coding genes (BSTOLATCC_MAC117 and BSTOLATCC_MAC118) are similar to this tRNA (Extended Data Fig. 1f; 59-71% identity to Contig_39-tRNA-like(UCA)). In the YAMAT-seq reads all four putative RNAs show signs of base modifications at a similar position, as well as non-templated "CCA" tail addition. Given that UCA anticodons may be the determinant of the ability of tryptophanyl-tRNA synthetases to charge tRNAs[23,24], the charging status of these molecules should be determined in future.

As judged by BLASTN searches with parameters for more sensitive short match detection ("-task blastn-short"), none of these tRNA-like molecules are similar to other *Blepharisma* tRNA genes. We also found homologs of these putative tRNAs in MAC genomes of other *Blepharisma* species by BLASTN searches, but not in *Stentor coeruleus*. Multiple sequence alignments of the region corresponding to the tRNA-like genes on *B. stoltei*'s Contig_39 with syntenic regions from two other *Blepharisma* species, show substantially more sequence conservation in these tRNA-like molecule genes than their flanking regions, i.e. 93.8% identity for the 72 bp region of the tRNAs vs. 60.5% and 61.4% identity in the flanking regions of the same length (Extended Data Fig. 1e). This suggests these genes might not be pseudogenes and are subject to some degree of purifying selection. On the other hand, the substantial divergences of the genes from *B. stoltei* Contig_1 from the one on Contig_39 suggest the former may be pseudogenes.

#### *Blepharisma stoltei* has a pair of moderately divergent tryptophanyl-tRNA synthetases

As pointed out in Kachale et al. 2023, the strict anticodon recognition capabilities of tryptophanyl-tRNA synthetases in eukaryotes like yeast and plants mean that a single synthetase may not be able to recognize both CCA and UCA anticodons, and thus, mutations outside of the anticodon may need to arise to permit efficient UGA translation.

*Blepharisma stoltei* has two non-mitochondrial tryptophanyl-tRNA synthetases (TrpRS1 — BSTOLATCC_MAC22805 and TrpRS2 — BSTOLATCC_MAC21555; 72% identity at the amino acid level), unlike any other ciliate species we examined (including *Stentor coeruleus*, *Fabrea salina* and *Spirostomum ambiguum*) with an assembled MAC genome. *Blepharisma*, like other model ciliates, also encodes a single mitochondrial tryptophan synthetase (mt-TrpRS) gene (BSTOLATCC_MAC5053). The most notable difference between TrpRS1 and TrpRS2 is that the latter has a 25 amino acid (aa) N-terminal peptide extension relative to the former. In the developmental RNA-seq time-series, TrpRS2 gene expression (average 3.7 RPKM for a developmental time series[25]) is a fraction of that of TrpRS1 (average 98.2 RPKM). Though we investigated the possibility the N-terminal peptide extensions of *Blepharisma* trpRS2 may be a signal peptide, none of the online prediction tools we used (e.g. Signalp 5.0[26]) predicted such a peptide. It is possible that the weaker expressed gene has a different function, but this, and its ability to charge *Blepharisma* $tRNA^{Trp}$'s will need to be experimentally ascertained.

Curiously, in BLASTP searches of the *Euplotes octocarinatus* genome database[27], we noticed this ciliate species, which uses a distinct $tRNA^{Cys}{}_{UCA}$ for UGA translation, has two distinct cysteine tRNA synthetases (82% amino acid identity): one of which has an N-terminal extension of 28 aa (Contig31990.g20673.t1) relative to the other (Contig16145.g5737.t1). Evolution of such paralogs raises the possibility of recognition of distinct tRNA species by each.

#### Bias of base downstream of *Blepharisma* stop codons and UGA tryptophan codons

In bacteria and eukaryotes, stop codon recognition is considered to involve additional bases flanking these triplets, with the strongest contribution from the downstream (+4) base[28]. In *Condylostoma magnum*, we observed some biases both downstream (+4) and upstream (-1) of the stop codons[1]. In *Blepharisma*, there may be a +4 bias in its less frequent UAG stops but not the more common UAA ones (Extended Data Fig 2d and e). The -1 base upstream of both *Blepharisma* stops may also be slightly biased. In *B. stoltei* the +4 base downstream of UGA is most often adenosine (52.1%; Extended Data Fig. 2b). UGA is also the codon with the highest adenosine frequency at this position (Extended Data Fig. 2c; no such bias occurs for UGG tryptophan codons; Extended Data Fig. 2f). Tryptophan incorporation in yeast using near-cognate tRNAs is favoured in translational readthrough of UGA with +4 adenosine[29]. The adenosine bias in *Blepharisma* may signal favouring of translation and avoidance of +1 bases that may promote premature termination by its eRF1's with some residual capacity to terminate at UGA codons, as suggested by previous *in vitro* experiments[9].

### Supplementary Methods

#### Cell cultivation and harvesting

For genomic DNA isolation *Blepharisma stoltei* ATCC 30299, *B. stoltei HT-IV*, *B. japonicum* R1072 and *B. undulans* K78 cells were cultured in Synthetic Medium for Blepharisma (SMB)[30] at 27 ℃. Blepharismas were fed *Chlorogonium elongatum* grown in Tris-acetate phosphate (TAP) medium[31] at room temperature. *Chlorogonium* cells were pelleted at 1500 g at room temperature for 3 minutes to remove most of the TAP medium, and resuspended in 50 mL SMB. 50 mL of dense *Chlorogonium* was used to feed 1 L of *Blepharisma* culture once every three days.

*Loxodes magnus* cells were cultured as previously described[11,32].

#### DNA isolation and sequencing

Total genomic DNA from *Blepharisma stoltei* ATCC 30299, *B. stoltei* HT-IV and *B. undulans* K78 was isolated with the SigmaAldrich GenElute Mammalian genomic DNA kit. A sequencing library was prepared with a NEBnext FS DNA Library Prep Kit for Illumina and sequenced on an Illumina HiSeq 3000 sequencer, generating 150 bp paired-end reads. Total genomic DNA from *B. japonicum* was isolated with the Sigma-Aldrich GenElute Mammalian genomic DNA kit and and sequencing library was prepared with the TruSeq Nano DNA Library Prep Kit (Illumina) and sequenced on an Illumina NovaSeq6000, generating 150 bp paired-end reads.

The *Loxodes magnus* DNA isolation procedure will be reported in future (Seah et al., manuscript in preparation).

#### Genomes and transcriptomes used

The following published ciliate macronuclear (MAC) genomes were used to search for tryptophan tRNAs: *Condylostoma magnum* - INSDC assembly accession GCA_001499635; *Blepharisma stoltei* ATCC30299 - https://bleph.ciliate.org/common/downloads/bleph/Bsto_ATCC_MAC_genome.fa; *Fabrea salina* - accession GCA_022984795; *Stentor coeruleus* - accession GCA_001970955.

For eRF1 sequences in Fig. 2a we used the following transcriptomes: a transcriptome of *Climacostomum virens* MMETSP1397; we assembled a *Plagiopyla frontata* transcriptome from RNA-seq deposited under BioProject PRJNA542330 using RNASpades[33] from the SPAdes[34] package version 3.14.0 with default parameters; aside from *Loxodes magnus*, karyorelict transcriptomes were previously assembled[2] and are available from EDMOND[35]. Aside from the same MAC genomes as those used for the tRNA searches, the following genomes were used for the eRF1 sequences in Fig. 2a: *Tetrahymena thermophila* MAC genome 2020 from ciliate.org; *Euplotes octocarinatus* from EOGD (http://ciliates.ihb.ac.cn/database/home/#eo); *Spirostomum ambiguum* PRJCA009852 (https://ngdc.cncb.ac.cn/). *Nyctotherus ovalis* eRF1 sequences are from GenBank accessions AY517526 and AY517527.

DNA-seq reads for *Blepharisma stoltei* HT-IV and *Blepharisma japonicum* were trimmed with BBDuk (version 38.22)[36], with the command and switches: "BBDukF -Xmx27g, threads=8, in=R1.gz, in2=R2.gz, ref=phix174_ill.ref.fa.gz,adapters.fa, qtrim=r, trimq=28, ktrim=r, k=31, mink=11, hdist=1, tpe, tbo". We assembled the macronuclear genomes of *Blepharisma stoltei* HT-IV, *B. japonicum* and *B. undulans* using SPAdes[34] (version 3.13) with kmers 21, 33, 55, 77, 99, and 127. These genomes were deposited in BioProject PRJEB60781, under accessions ERZ16477024, ERZ16477415 and ERZ16477695, respectively.

The *Loxodes magnus* genome assembly procedure will be reported in future (Seah et al., manuscript in preparation).

The *Blepharisma stoltei* ATCC30299 mitochondrial genome is available from accession ERZ16464618.

#### tRNA prediction

tRNA genes were predicted using tRNAscan-SE 2.0[18] with default parameters for eukaryotes and ARAGORN v1.2.38[19] with default parameters. tRNA secondary structures shown in Fig. 1 are based on visual inspection of both ARAGORN and tRNAscan-SE 2.0 predictions. The discriminator base just prior to the tRNA CCA tails was judged to be a purine for *Fabrea salina* based on experimental characterization of discriminator base preferences[37].

One tRNA with a UCA codon was predicted by tRNAscan-SE in the *L. magnus* MAC genome. However, this prediction had a low score 31.0 with an unpaired base in one of the stems that led to a pseudogene classification, and no YAMAT-seq reads were mapped to its genes. tRNAscan-SE predicted 332 pseudogenes out of a total of 9051 tRNA genes in this genome.

The minimum free energy tRNA secondary structure used to generate Extended Data Fig. 1d was predicted using the RNAfold[20] web server with default parameters.

#### tRNA sequence library preparation and processing

Total RNA was extracted from the ciliate cells using TRI Reagent (Sigma) according to the manufacturer's protocol, and treated with DNase. This RNA was used to produce a library enriched in mature tRNAs by the YAMAT-seq method[38]. This method captures mature tRNAs by using a Y-shaped adapter complementary to the CCA-3' tails synthesised during tRNA maturation. The resulting multiplexed cDNA library was sequenced on either an Illumina MiSeq, HiSeq 3000 or NextSeq sequencer in paired-end mode. YAMAT-seq reads for *B. stoltei* ATCC30299 and *Loxodes magnus* LM5 are available from BioProject PRJEB60781, accessions ERR11079929 and ERR11079930, respectively.
Paired-end YAMAT-seq reads were merged with BBMerge (default parameters) from the BBTools package (version 37.62)[39]. Forward and reverse adapters were trimmed off the merged reads using cutadapt 3.2[40] (default parameters; -m 20). Using tRAX (cloned from https://github.com/UCSC-LoweLab/tRAX on 12 March 2021)[41] with default parameters, YAMAT-seq reads were mapped to

tRNAs predicted by tRNAscan-SE 2.0, for read quantification and inspection of potential base modifications.

Of the ~34 million unique reads mapped to the predicted *B. stoltei* tRNAs ("samtools view -F 0x904 -c"), ~173,000 mapped to the 6 identical $tRNA^{Trp}_{CCA}$ *B. stoltei* genes. Of the ~21 million reads mapped to the *L. magnus* MAC genome, ~14,200 mapped to $tRNA^{Trp}_{CCA}$ genes (Supplementary Table 1; Source Data Fig. 1). Reverse transcription error-rates are very low, on the order of $10^{-6}$ for C to U[42]. Thus, C to U anticodon base editing should be distinguishable if it occurs at a higher frequency. Clear signals of modified bases were present in the reads mapped to $tRNA^{Trp}_{CCA}$ genes of both organisms at non-anticodon positions.

To examine mapping of YAMAT-seq reads to potential tRNAs which tRNAscan-SE 2.0 did not predict, we mapped the same reads to the *B. stoltei* ATCC30299 MAC and mitochondrial genomes using a version of HISAT2[22] version 2.0.0-beta, modified to permit shorter introns, with parameters --min-intronlen 9 --max-intronlen 30. This version of HISAT2 has the static variable minIntronLen in hisat2.cpp lowered to 9 from 20 (available from a HISAT2 GitHub fork: https://github.com/Swart-lab/hisat2/; commit hash 86527b9).

#### Multiple sequence alignment and phylogenetic inference

MAFFT version 7.450 (parameters: algorithm FFT-NS-2, BLOSUM62 scoring matrix; gap opening penalty 1.53; offset value 0.123)[43,44] was used to align the eRF1 sequences used in Fig. 2a. RAxML version 8.2.11 was used to infer the phylogeny in the same figure, using the switches "-m PROTGAMMAJTT -f d -N 1 -p 1".

Though we did not analyse them, 1:1 orthologs of the eRF1 paralogs in *B. stoltei* are also present in the *B. japonicum* MAC genome we assembled, with the same set of position 67 amino acids (i.e. one with Ala67, two with Ser67).

To search for similar tRNAs to the tRNA-like molecule, Contig_39-tRNA-like(UCA) from *B. stoltei* ATCC30299 in the genomes of *B. stoltei* HT-IV, *B. japonicum* R1072 and *B. undulans* K78 we used BLASTN with the switch "-task blastn-short". MAFFT version 7.450 (parameters: algorithm E-INS-i, 200PAM/k=2 scoring matrix; gap opening penalty 1.53) was used to align these sequences flanked by 80 bp of surrounding sequences (Extended Data Fig. 1e). MAFFT version 7.450 with the same parameters was used to align the *B. stoltei* ATCC30299 paralogs in Extended Data Fig. 1f.

### Supplementary Tables

#### Supplementary Table 1

Anticodon editing given for tRNAs with YAMAT-seq reads mapped to them. Note that two *L. magnus* tRNAs (tRNA-Trp-CCA-000147F and tRNA-Trp-CCA-7822) have no reads mapped to them, with the former a likely pseudogene (potential mutant of $tRNA^{Gln}_{CTA}$) and the latter also unusual and identified by tRNAscanSE and not ARAGORN, and thus neither are included in the table. “Structure” and “Variant” indicate tRNAs that either appear distinct or highly similar (few substitution differences), respectively.

| | | | | **Anticodon base 1 counts** | | | | | **Anticodon base 2 counts** | | | | | | |
|---|---|---|---|---|---|---|---|---|---|---|---|---|---|---|---|
| **Organism** | **AS stem** | **tRAX identifier** | **%C** | **C** | **A** | **T** | **G** | **%C** | **C** | **A** | **T** | **G** | **Structure** | **Variant** | **Pseudogene** |
| *B. stoltei* | 4 bp | tRNA-Trp-CCA-12 | 99.96 | 172863 | 22 | 41 | 3 | 99.95 | 172861 | 34 | 57 | 2 | 1 | 1 | No |
| *L. magnus* | 4 bp | tRNA-Trp-CCA-000066F | 99.71 | 3109 | 9 | 0 | 0 | 99.84 | 3114 | 5 | 0 | 0 | 1 | 1 | No |
| *L. magnus* | 4 bp | tRNA-Trp-CCA-000147F | 99.86 | 6606 | 9 | 0 | 0 | 99.92 | 6610 | 5 | 0 | 0 | 1 | 2 | No |
| *L. magnus* | 4 bp | tRNA-Trp-CCA-000518F | 99.71 | 3108 | 9 | 0 | 0 | 99.84 | 3113 | 5 | 0 | 0 | 1 | 1 | No |
| *L. magnus* | 5 bp | tRNA-Trp-CCA-000035F | 98.91 | 543 | 0 | 6 | 0 | 98.72 | 542 | 0 | 7 | 0 | 3 | 1 | Likely |
| *L. magnus* | 5 bp | tRNA-Trp-CCA-000109F | 99.76 | 830 | 2 | 0 | 0 | 98.32 | 819 | 1 | 5 | 8 | 2 | 1 | Likely |

#### Supplementary Table 2

Incorrect genetic code assignments (non-exhaustive) for ciliates in the NCBI Taxonomy database (accessed 17 March 2023). Taxa are generally assumed to have the same genetic code as their parent taxon unless there is evidence to the contrary. Due to the frequency of genetic code changes in ciliates, genetic code annotations have to be revised more carefully for this clade.

| Name | Taxid | Rank | NCBI code | Correct code | References | Notes |
|---|---|---|---|---|---|---|
| *Fabrea* | 342562 | genus | 10 | 1 | Heaphy et al. 2017, Swart et al. 2017 | |
| *Climacostomum* | 49979 | genus | 10 | 1 | Heaphy et al. 2017, Swart et al. 2017 | |
| *Stentor* | 5962 | genus | 10 | 1 | Slabodnick et al. 2017 | At least *Stentor coeruleus*, likely others too |
| *Blepharisma* | 5959 | genus | 4 | NA | Heaphy et al. 2017, Swart et al. 2017; Singh et al. 2023; present study | The deprecated and removed "Blepharisma genetic code" 15 also contains incorrect assignment for UAG |
| Karyorelictea | 33827 | class | 6 | 27 | Seah et al. 2022 | |
| *Nyctotherus* | 70074 | genus | 6 | 1 | Ricard et al. 2008 | |
| *Nyctotherus ovalis* | 70075 | species | 10 | 1 | Ricard et al. 2008 | Other Metopidae also likely to have code 1 (Yan et al. 2019) |
| Peritrichia | 6021 | class | 6 | 30 | Sánchez-Silva 2003 | |
| *Plagiopyla frontata* | 35117 | species | 1 | 27 | McGowan et al. 2022 | |
