## Supplementary material for "How did UGA codon translation as tryptophan evolve in certain ciliates? A critique of Kachale et al. 2023 *Nature*": Source Data: README.txt

This archive contains:
1. tRNA predictions from ARAGORN (.aragorn) and tRNAscan-SE 2.0
(.tRNAscanSE.), for the MAC and mitochondrial genomes of the ciliates
Blepharisma stoltei ATCC30299 (Bsto), Blepharisma japonicum R1072, Blepharisma
undulans K78, and Loxodes magnus LM5 (Lmag). tRNA predictions are also
provided for Paramecium biaurelia from ParameciumDB
(https://paramecium.i2bc.paris-saclay.fr/).
2. SAM files containing YAMAT-seq reads mapped by tRAX to Blepharisma stoltei
and Loxodes magnus MAC genome-encoded Trp-tRNAs. Also included are the
reference sequences to which tRAX mapped the reads (.fa files). 
